# Heterogeneity in slow synaptic transmission diversifies Purkinje cell timing

**DOI:** 10.1101/2024.03.24.585592

**Authors:** Riya Elizabeth Thomas, Franziska Mudlaff, Kyra Schweers, W. Todd Farmer, Aparna Suvrathan

**Affiliations:** Centre for Research in Neuroscience, Brain Repair and Integrative Neuroscience Program, Research Institute of the McGill University Health Centre, Montréal, Québec H3G 1A4, Canada; Integrated Program in Neuroscience, McGill University, Montréal, Québec H3G 1A4, Canada; McGill University, Montréal, Québec H3G 1A4, Canada; Department of Pharmacology, University of North Carolina at Chapel Hill, Chapel Hill, NC 27599, USA, Neuroscience Center, University of North Carolina at Chapel Hill, Chapel Hill, NC 27599, USA; Department of Neurology and Neurosurgery, McGill University, Montréal, Québec H3G 1A4, Canada; Department of Pediatrics, McGill University, Montréal, Québec H3G 1A4, Canada

## Abstract

The cerebellum plays an important role in diverse brain functions, ranging from motor learning to cognition. Recent studies have suggested that molecular and cellular heterogeneity within cerebellar lobules contributes to functional differences across the cerebellum. However, the specific relationship between molecular and cellular heterogeneity and diverse functional outputs of different regions of the cerebellum remains unclear. Here, we describe a previously unappreciated form of synaptic heterogeneity at parallel fiber synapses to Purkinje cells. In contrast to uniform fast synaptic transmission, we found that the properties of slow synaptic transmission varied by up to three-fold across different lobules of the mouse cerebellum, resulting in surprising heterogeneity. Depending on the location of a Purkinje cell, the time of peak of slow synaptic currents varied by hundreds of milliseconds. The duration and decay-time of these currents also spanned hundreds of milliseconds, based on lobule. We found that, as a consequence of the heterogeneous synaptic dynamics, the same brief stimulus was transformed into prolonged firing patterns over a range of timescales that depended on Purkinje cell location

## Introduction

The cerebellum supports a number of functions, far beyond the motor domain (Apps and Hawkes, 2009; Stoodley, Valera and Schmahmann, 2012; Witter and De Zeeuw, 2015; Apps *et al*., 2018; De Zeeuw, Lisberger and Raymond, 2021). Yet, cerebellar cortical microcircuits are known for their uniformity, consisting of repeated stereotyped modules (Ruigrok, 2011; Cerminara *et al*., 2015; Valera *et al*., 2016). The homogeneity of cerebellar architecture has led to the assumption that the computations performed are similar across regions (Schmahmann, 2010), in spite of their diverse functional roles. However, more recent discoveries demonstrated a marked degree of molecular and cellular heterogeneity across cerebellar lobules (Ebner *et al*., 2012; Zhou *et al*., 2014, 2015; Cerminara *et al*., 2015; Tsutsumi *et al*., 2015; Witter and De Zeeuw, 2015; Nguyen-Minh *et al*., 2019). The functional impact of this heterogeneity is not well understood. The properties of Purkinje cells, the output neurons of the cerebellar cortex, vary across regions, e.g. with respect to dendritic integration (Eccles, Ito and Szentagothai, 1967), firing patterns (Zhou *et al*., 2014, 2015), and axonal output (Voogd, 2011). However, potential heterogeneity of Purkinje cells inputs is not well explored.

The inputs to Purkinje cells are well known to carry diverse information ((Bower and Woolston, 1983; Brodal and Bjaalie, 1997). For instance, parallel fibers (PFs) can encode sensory (Bosman *et al*., 2010; Shimuta, Sugihara and Ishikawa, 2020), motor (Wiestler, McGonigle and Diedrichsen, 2011; Proville *et al*., 2014), or reward (Wagner *et al*., 2017) features. Thus, there is reason to hypothesize that there is heterogeneity at these inputs. The glycolytic enzyme aldolase C or zebrin II is a well-investigated marker of Purkinje cell heterogeneity, which patterns the cerebellar cortex into parasagittal zebra-like bands. Zebrin bands can receive similar input information and send similar output information (Sugihara and Shinoda, 2004; Voogd and Ruigrok, 2004; Pijpers *et al*., 2006; Apps and Hawkes, 2009; Ruigrok, 2011), although functional units within the cerebellum can also cross zebrin boundaries (Graham and Wylie, 2012). Zebrin identity correlates with some forms of molecular heterogeneity, including components of the metabotropic glutamate receptor 1 (mGluR1) signaling cascade (Mateos *et al*., 2001; Wadiche and Jahr, 2005; Furutama *et al*., 2010; Wu *et al*., 2019).

At PF to Purkinje cell synapses, excitatory synaptic transmission is composed of a fast, α-amino-3-hydroxy-5 methyl-4-isoxazolepropionic acid receptor (AMPAR)-dependent input, as well as a slower mGluR1-dependent component (Batchelor, Madge and Garthwaite, 1994; Batchelor and Garthwaite, 1997; Tempia *et al*., 1998). The metabotropic arm of PF input triggers an intracellular signaling cascade, and a slow excitatory postsynaptic current (slow EPSC) through the transient receptor potential channel TRPC3 (Hartmann *et al*., 2008; Hartmann, Henning and Konnerth, 2011; Henning, 2011; Hartmann and Konnerth, 2015). mGluR1 signaling (Aiba *et al*., 1994; Ichise *et al*., 2000; Hartmann *et al*., 2008; Lüscher and Huber, 2010; Becker, 2014; Crupi, Impellizzeri and Cuzzocrea, 2019; Cole and Becker, 2023) and TRPC3-dependent currents (Hartmann *et al*., 2008; Hartmann, Henning and Konnerth, 2011; Cole and Becker, 2023) are both essential for normal cerebellar function. However, heterogeneity of mGluR1-mediated slow synaptic transmission across lobules has never yet been investigated.

Here, we tested the hypothesis that slow synaptic transmission at PF synapses varies across different lobules of the cerebellum. To do so, we measured synaptic currents at PF inputs to Purkinje cells in acute mouse brain slices. We found that properties of slow synaptic currents varied up to three-fold across different cerebellar lobules. As a consequence, PF inputs were transformed into a diverse range of prolonged firing outputs that depended on Purkinje cell location. This previously-unappreciated heterogeneity in slow synaptic transmission thus diversifies Purkinje cell firing dynamics.

## Results

### PF-driven slow EPSCs were heterogeneous across cerebellar lobules

A remarkable feature of the cerebellar cortex is its stereotypical anatomy, composed of well-defined regions and lobules that can be reproducibly identified. However, it remains unclear whether Purkinje cells in different cerebellar regions and lobules show uniform or distinct synaptic properties, and how this may contribute to region or lobule-specific cerebellar cortex output. To help resolve this, we compared the physiological properties of slow EPSCs at Purkinje cells in lobules located in the midline vermis with those in the laterally positioned floccular lobule (Figure 1a), regions that have different functional roles (Cerminara *et al*., 2015; Witter and De Zeeuw, 2015) and molecular profiles (Fujita *et al*., 2014). We measured PF driven slow EPSCs using patch-clamp recordings from Purkinje cells (see Methods) in different lobules of the cerebellum taken from acute mouse brain slices (Figure 1b). The slow EPSC at PF to Purkinje cell synapses is measured in response to physiologically relevant high-frequency PF stimulation in the presence of antagonists of AMPARs and GABA_A_ (γ-amino butyric acid_A_) receptors (Batchelor and Garthwaite, 1997; Finch and Augustine, 1998; Brasnjo and Otis, 2001; Isope and Barbour, 2002; Canepari, Auger and Ogden, 2004; Chadderton, Margrie and Häusser, 2004; Jorntell and Ekerot, 2006; Hartmann *et al*., 2008; van Beugen *et al*., 2013).

**Figure 1.**
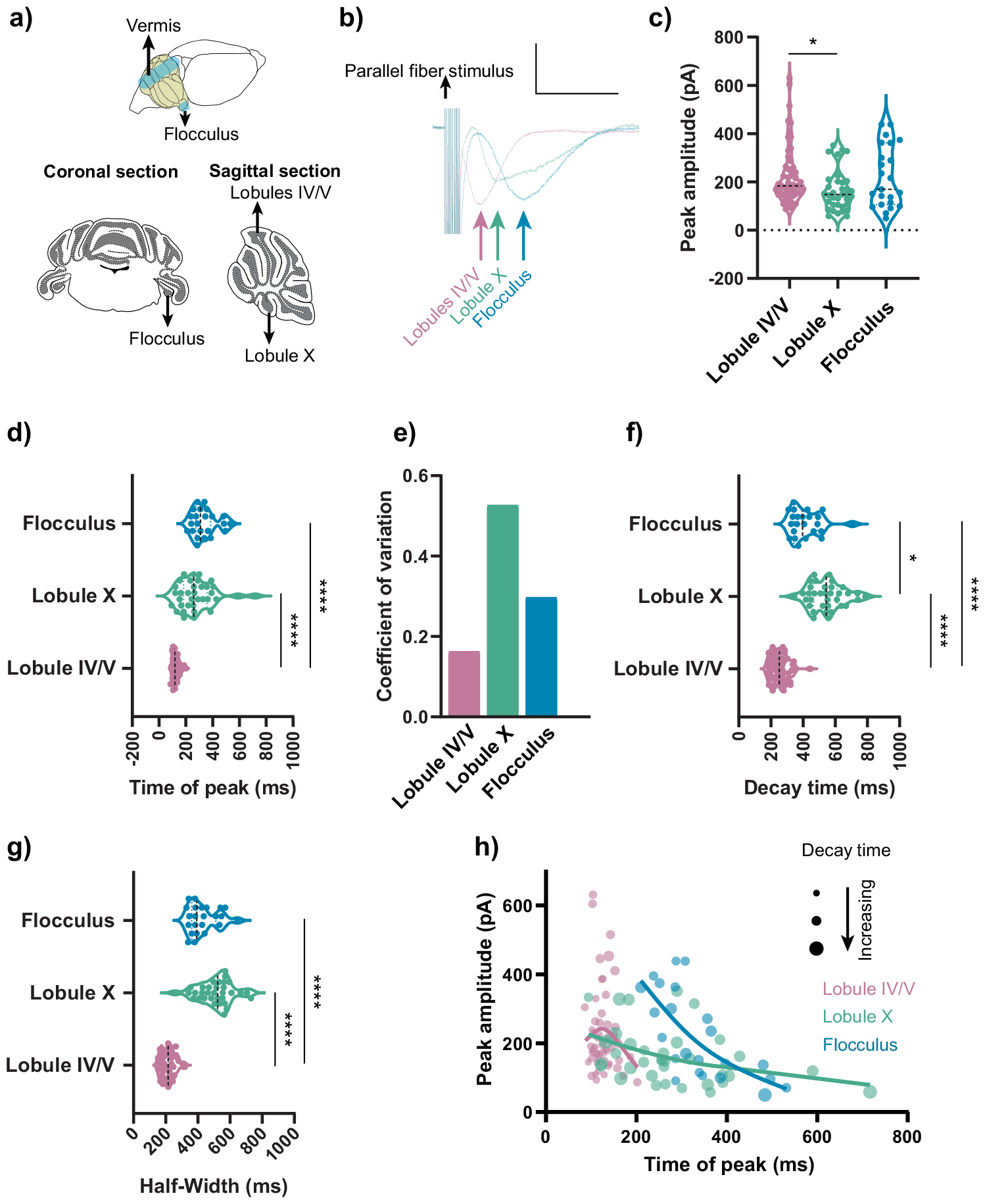
The slow EPSC (sEPSC) was heterogenous across cerebellar lobules. a) Schematic representation of a mouse brain highlighting the recording regions: the cerebellar flocculus and vermis. Parasagittal acute cerebellar slices for recordings in vermis lobule IV/V and lobule X and coronal sections for flocculus recordings. b) Representative sEPSC traces show delayed peak time in lobule X and the flocculus compared to lobule IV/V. Scale bar: 100 pA, 500 ms. c) Peak amplitudes of sEPSCs were similar across lobules. (*p<0.05). d) sEPSC peak time was earlier in lobule IV/V in comparison to lobule X and the flocculus. (****p<0.0001). e) High heterogeneity in the time of peak within lobules X and flocculus compared to lobule IV/V is demonstrated by higher coefficients of variation. f) Lobule X sEPSCs exhibited the longest 90% to 10% decay time. Both lobule X and flocculus sEPSCs had longer decay times than those in lobule IV/V. (*p<0.05, ****p<0.0001). g) A higher half-width sEPSC was observed in lobule X and the flocculus compared to lobule IV/V. (****p<0.0001). h) sEPSCs with similar peak amplitudes in lobule IV/V, lobule X and flocculus had different times of peak. Bubble size represents the decay time of the sEPSC. Data from each lobule was fit with a spline in equivalent color. Statistical significance was assessed using Kruskal-Wallis analysis of variance on ranks followed by Dunn’s multiple comparisons test. Violin plots show median and quartiles.

We found that slow EPSCs in vermal lobule IV/V had dynamics surprisingly different to those in the flocculus (Figure 1b, c, d). In particular, the time of peak of lobule IV/V slow EPSCs was up to three-fold lower than the time of peak of floccular slow EPSCs (Figure 1d, e). Moreover, slow EPSCs in lobule IV/V were less variable than those in the flocculus (Figure 1d, e). Next, we measured slow EPSCs from lobule X of the vermis. Lobule X is similar to the flocculus in terms of being largely zebrin-positive, in contrast to the largely zebrin-negative lobule IV/V. We found that the times of peak of slow EPSCs in lobule X were similar to those in the flocculus, and more delayed than those in lobule IV/V. Purkinje cells in lobule X, like those in the flocculus, displayed heterogeneous timing of their slow current events (Figure 1d). The heterogeneity we observed in slow EPSC dynamics was also reflected by slower decay times and longer half-widths in vermal lobule X and the flocculus, compared to vermal lobule IV/V (Figure 1f, g). In addition, the slow EPSC had a slightly lower amplitude in lobule X, which aligned with previous findings (Figure 1c) (Wadiche and Jahr, 2005). Next, we measured how the size of the EPSC impacts its timing by comparing the peak amplitude vs. time of peak of each cell across the three lobules. Regardless of the peak amplitude of the slow EPSC in lobule IV/V, the time of peak had a low variance when compared to other lobules (pink, Figure 1h). In marked contrast, slow EPSCs from the flocculus (blue, Figure 1h) formed a non-overlapping population with a delayed time of peak. Finally, slow EPSCs from lobule X (green) showed a time of peak in between these two populations but were distinguished by their longer decay times. Thus, Purkinje cells possess distinct timing and shape of slow EPSCs depending on their location in different lobules.

In contrast to the differences observed with the slow ESPCs, we could not detect a difference in the timing of fast EPSCs in lobule IV/V and flocculus (Figure S1). Thus, the timing heterogeneity in slow EPSCs was specific to this current and was not a feature of the synapse or the cell.

### Zebrin identity did not determine slow EPSC properties

The differences in slow EPSCs in lobule IV/V and flocculus could be due to a variety of molecular and cellular signatures of Purkinje cells in these lobules. One molecular signature that distinguishes Purkinje cells with different properties is zebrin. It was previously shown that Purkinje cells in vermal lobule IV/V are largely zebrin-negative while those in vermal lobule X and the flocculus are largely zebrin-positive (Fujita *et al*., 2012) (Figure 2a, b, c, d). However, some Purkinje cells within these lobules do not follow the same zebrin identity as the majority of other Purkinje cells within the lobule. We took advantage of this intrinsic variability to interrogate how the zebrin identity of individual Purkinje cells relates to their slow EPSC timing. In order to do this, patch-clamp recordings from different regions were first performed, during which the recorded cell was dye filled. Slices were subsequently fixed and immunolabelled retrospectively for zebrin (Figure 2e, f). This way, the zebrin identity of individual cells was identified after characterizing its slow EPSC. As expected, most cells in each region followed the zebrin identity of that lobule. However, as expected, instances were found where Purkinje cells had the opposite zebrin identity (Figure 2g). Surprisingly, we found that the identity of the lobule rather than an individual Purkinje cell’s zebrin identity defined the timing of the slow EPSC (Figure 2g). For example, the sparse zebrin-positive Purkinje cells in lobule IV/V had slow EPSC currents that followed the relatively fast kinetics of the majority zebrin-negative cells within that region. A mirror effect was seen for Purkinje cells in the flocculus and lobule X, where sparse zebrin-negative cells showed slow EPSCs with kinetics similar to Purkinje cells that were zebrin-positive. Thus, our results show that cerebellar lobule rather than individual Purkinje cell zebrin expression determined slow EPSC properties.

**Figure 2.**
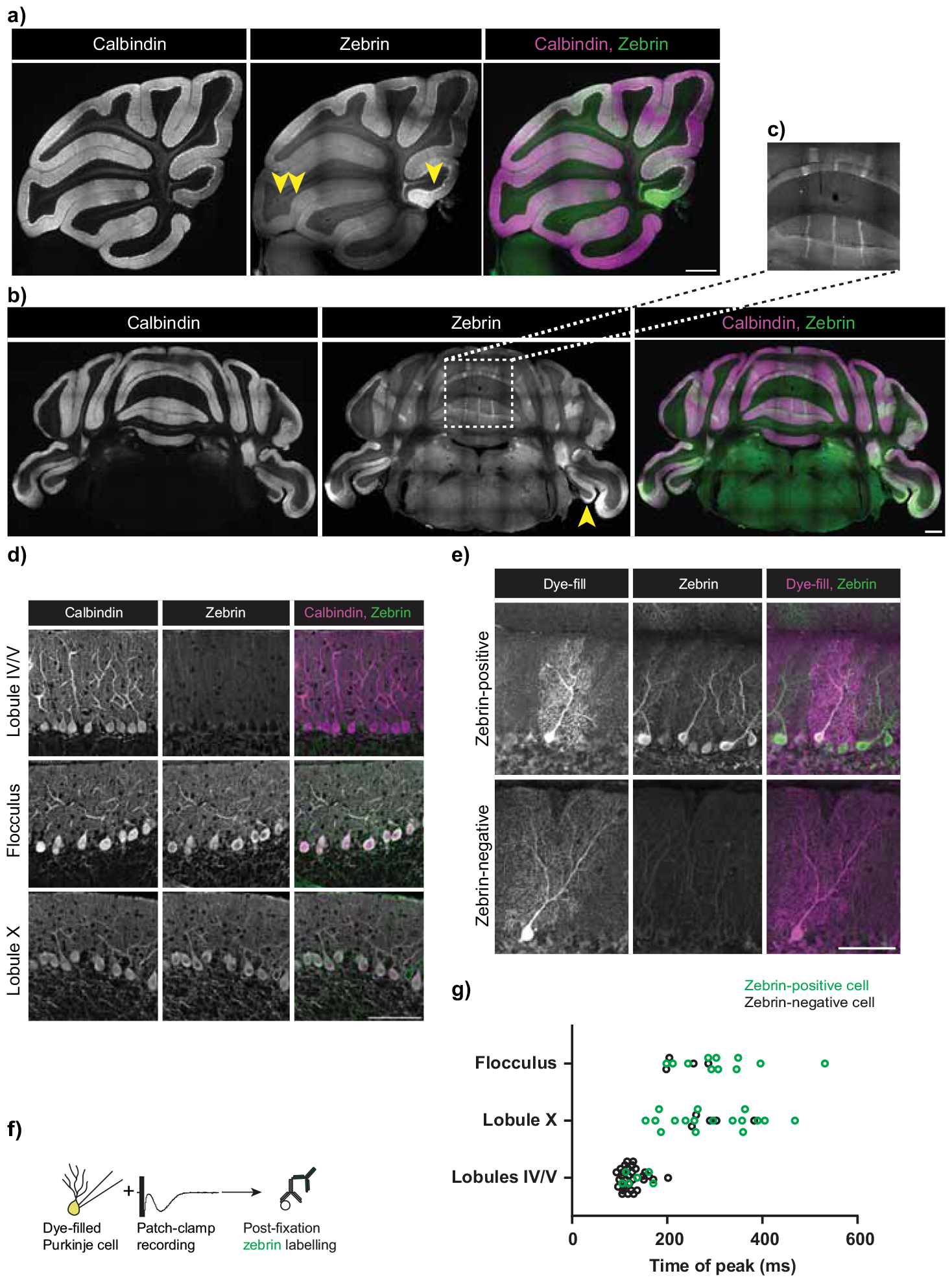
Zebrin/Aldolase C expression did not predict sEPSC peak latency. a) Stitched mouse vermis sagittal sections, immunolabelled with anti-calbindin (which marks Purkinje cells) and anti-zebrin antibodies. Rightmost panel: Merged. Single arrowhead marks lobule X, double arrowhead marks lobule IV/V. Scale bar: 500 μm b) Stitched coronal sections of the mouse cerebellum, immunolabelled with anti-calbindin and anti-zebrin antibodies. Rightmost panel: Merged. Single arrowhead marks flocculus. Scale bar: 500 μm. c) Inset illustrates zebrin stripes. d) Zebrin-negative Purkinje cells in lobule IV/V, and zebrin-positive Purkinje cells in the flocculus and lobule X. Scale bar: 100 μm. e) Representative image of a 300 μm thick cerebellar section with an Alexa 488 dye-filled Purkinje cell, with the section retrospectively stained with anti-zebrin antibody. Upper row shows a dye-filled cell that is zebrin-positive, and lower row shows one that is a zebrin-negative. Scale bar: 100 μm. e) Schematic representing dye-fill of the whole-cell patch-clamped Purkinje cell from which sEPSCs were measured, followed by immunolabelling with anti-zebrin antibody. g) Zebrin identity did not predict time of peak of sEPSC, across regions. Most dye-filled lobule IV/V cells were zebrin-negative (black hollow circles) and most dye-filled lobule X and flocculus cells were zebrin-positive (green hollow circles). However, even zebrin positive cells (green hollow circles) in lobule IV/V had a characteristically shorter time of peak and zebrin-negative cells (black hollow circles) in lobule X and flocculus had a delayed time of peak.

### Slow EPSC heterogeneity persisted across a range of parameters

It is possible that the heterogeneity we observed is only present with the specific PF stimulation parameters we used (10 pulses at 100 Hz), which would limit its relevance. We ruled out this possibility by testing a range of parameters. We found that the heterogeneity in slow ESPCs remained even with a lower number of stimuli (starting with 3 pulses, shown to be the lowest number to activate mGluR1 (Brasnjo and Otis, 2001), and going up to 5 stimuli), (Figure 3a-i). In addition, we maintained the number of stimuli at 10 but tested a range of stimulation frequencies, from 50Hz to 200Hz. Across all frequencies, the time of peak of the slow EPSC in lobule IV/V was early and highly homogeneous, in contrast to the delayed and varied time of peak in lobule X and the flocculus. In all cases, the decay was different across lobules in a manner similar to that described in Figure 1 (Figure 4a-c). Thus, the specific number of stimuli or the frequency did not determine the heterogeneity in slow EPSCs.

**Figure 3:**
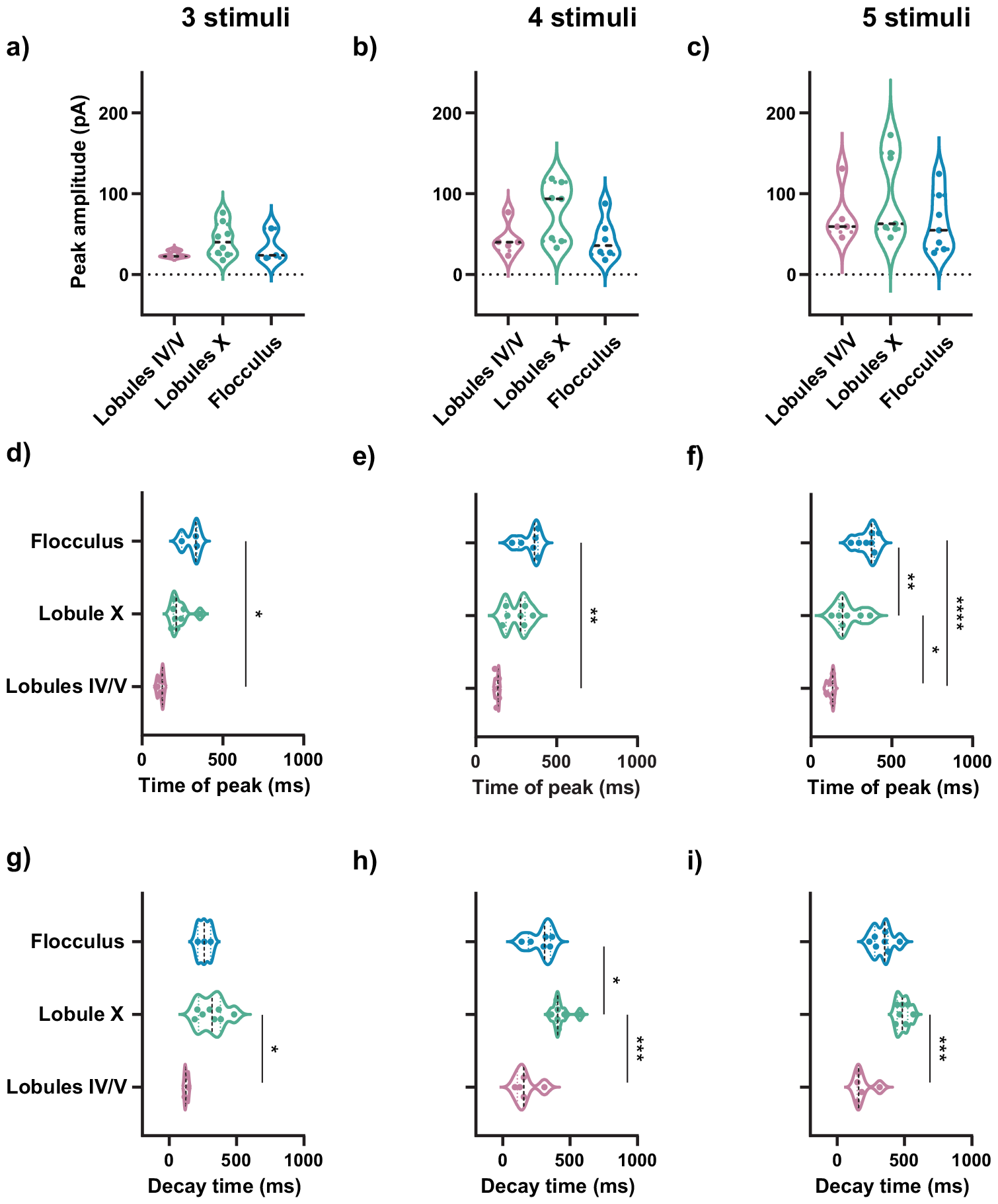
sEPSC heterogeneity did not depend on number of PF stimulations. (a,b,c) Peak amplitudes of sEPCS were similar in lobule IV/V, lobule X and flocculus in response to 3, 4, 5 PF stimulations respectively at 100Hz. (d,e,f) sEPSC time of peak was consistently earlier in lobule IV/V in comparison to lobule X and the flocculus, in response to 3, 4, 5 PF stimulations respectively at 100Hz. (g,h,i) sEPSC decay time was shorter in lobule IV/V in comparison to lobule X, and the flocculus fell in between the two, in response to 3, 4, 5 PF stimulations respectively at 100Hz. Statistical comparisons: (f,g,h) *p<0.05, **p<0.01, ***p<0.001,****p<0.0001, ordinary one-way ANOVA followed by Tukey’s multiple comparison test. (d,e,i)*p<0.05,**p<0.01, ***p<0.001, Kruskal-Wallis test followed by Dunn’s multiple comparison test. All data are Mean ± SEM. Violin plots show median and quartiles.

**Figure 4:**
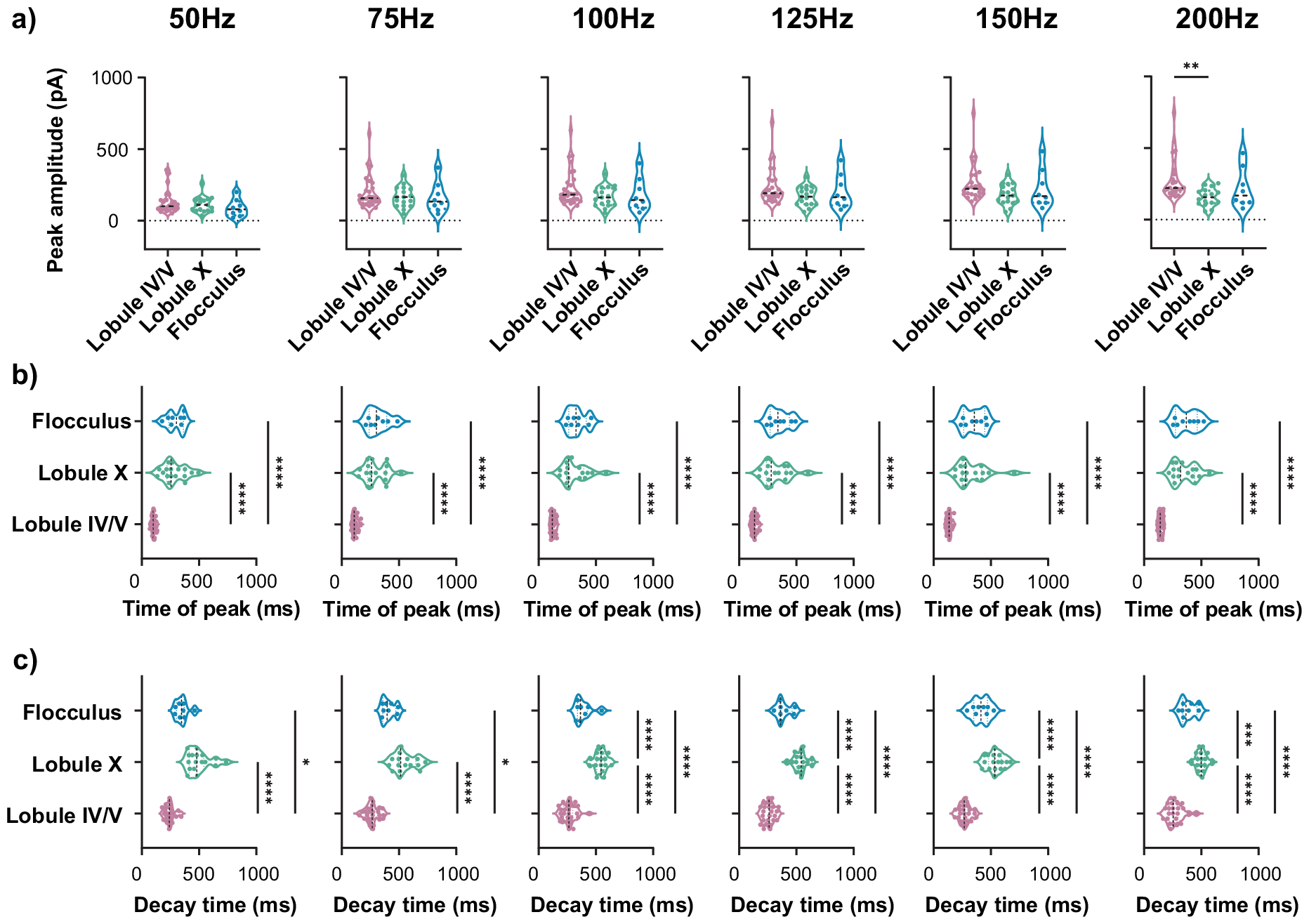
Heterogeneity of sEPSC properties persisted across a range of PF stimulation frequencies. (a) From left to right: Peak amplitude of sEPCS in lobule IV/V, lobule X and flocculus were similar, in response to 10 PF stimulations at 50,75,100, 125, 150 and 200Hz. **p<0.01, Kruskal-Wallis test followed by Dunn’s multiple comparison test. b) From left to right: sEPSC peak timing was shorter in lobule IV/V in comparison to lobule X and the flocculus, in response to 10 PF stimulations at 50, 75, 100, 125, 150 and 200Hz. ****p<0.0001, Kruskal-Wallis test followed by Dunn’s multiple comparison test. c) From left to right: Time for the sEPSC peak amplitude to decay from 90% to 10% was longest in lobule X, and was longer both in lobule X and the flocculus in comparison to lobule IV/V, in response to 10 PF stimulations at 50, 75, 100, 125, 150, 200Hz. 50 Hz, 75 Hz *p<0.05, ****p<0.0001, Kruskal-Wallis test followed by Dunn’s multiple comparison test. 100-200 Hz ***p<0.001,****p<0.0001, ordinary one-way ANOVA followed by Tukey’s multiple comparison test. Violin plots show medians and quartiles.

We were concerned that the time after break-in for whole cell patching could contribute to dialysis of the intracellular cytosol and impact pathways necessary for the slow EPSC (Sakmann and Neher, 1984; Rae and Fernandez, 1988; Cahalan and Neher, 1992). To account for this, we analyzed a subset of Purkinje cells from Figure 1 in which slow EPSCs were measured within the restricted time window of 10-20 min after break-in. The results showed that even within this restricted subset of cells, the heterogeneity observed across lobules remained (Figure S2).

Recent studies describe sex-based differences in cerebellar structure and physiology (Mercer *et al*., 2016; Steele and Chakravarty, 2018). To test this possibility, we compared slow EPSCs from male and female mice. We found no difference between slow EPSC properties of Purkinje cells from male and female mice (Figure S3).

### Heterogeneity in mGluR1-TRPC3 signaling may contribute to Purkinje cell slow EPSCs

The slow EPSC was previously shown to depend on mGluR1 (Hartmann *et al*., 2008). We wondered if the heterogeneity of the slow EPSC that we found relied on other types of signaling across the different lobules. However, we found that the non-competitive mGluR1 antagonist CPCCOEt (Hartmann *et al*., 2008) blocked slow EPSCs in all three regions (Figure S4a), arguing against this possibility.

Downstream of mGluR1, the slow EPSC is mediated by the non-selective cation channel TRPC3 (Hartmann *et al*., 2008; Chae *et al*., 2012; Cole and Becker, 2023). To test how TRPC3 blockade impacts the time of peak, we used the tyrosine kinase inhibitor genistein (Vazquez *et al*., 2004; Y. Kim *et al*., 2012), and the TRPC channel blocker SKF96365 (Chae *et al*., 2012; Song, Chen and Yu, 2014; Cole and Becker, 2023), which both partially block the slow EPSC, allowing us to measure the kinetics of the remaining current. We found that the magnitude of the slow EPSC was consistently reduced by genistein and by SKF96365 across lobules, but there was no shift in timing that could explain the differences between lobules (Figure S4b-e).

The characteristically prolonged time-course of the slow EPSC is related to the associated G-protein dependent signaling cascade (Figure 5a) (Brasnjo and Otis, 2001; Hartmann, Henning and Konnerth, 2011; Kano and Watanabe, 2017; Hirai and Kano, 2018; Cole and Becker, 2023). However, the signaling pathways connecting mGluR1 and TRPC3 signaling, and its regulation, are not fully understood (Cole and Becker, 2023). In spite of this, we wanted to identify candidate molecular players underlying the diversity of slow EPSCs across lobules. Therefore, we performed a bioinformatic analysis of Purkinje-cell transcriptomes using a published single-nucleus RNA sequencing dataset (Kozareva *et al*., 2021). We separated Purkinje-cell transcriptomes by lobule of interest and focused on molecular candidates related to the mGluR1-TRPC3 pathway (Figure 5b) (Brasnjo and Otis, 2001; Hartmann, Henning and Konnerth, 2011; Kano and Watanabe, 2017; Hirai and Kano, 2018; Cole and Becker, 2023). Comparison of gene expression demonstrated statistically significant differences across lobules. Key features of our analysis aligned with previous findings. For instance, expression of Grm1, which encodes mGluR1, was greater in lobule IV/V than in the flocculus (F) and lobule X. This is consistent with greater mGluR1b in zebrin-negative zones (Mateos *et al*., 2001). TRPC3 expression was also greater in the largely zebrin-negative lobule IV/V, in contrast to the largely zebrin-positive flocculus and lobule X (Wu *et al*., 2019). This is consistent with previous studies, although it is also known that TRPC3 expression does not perfectly overlap with zebrin zones (Blot *et al*., 2023).

**Figure 5.**
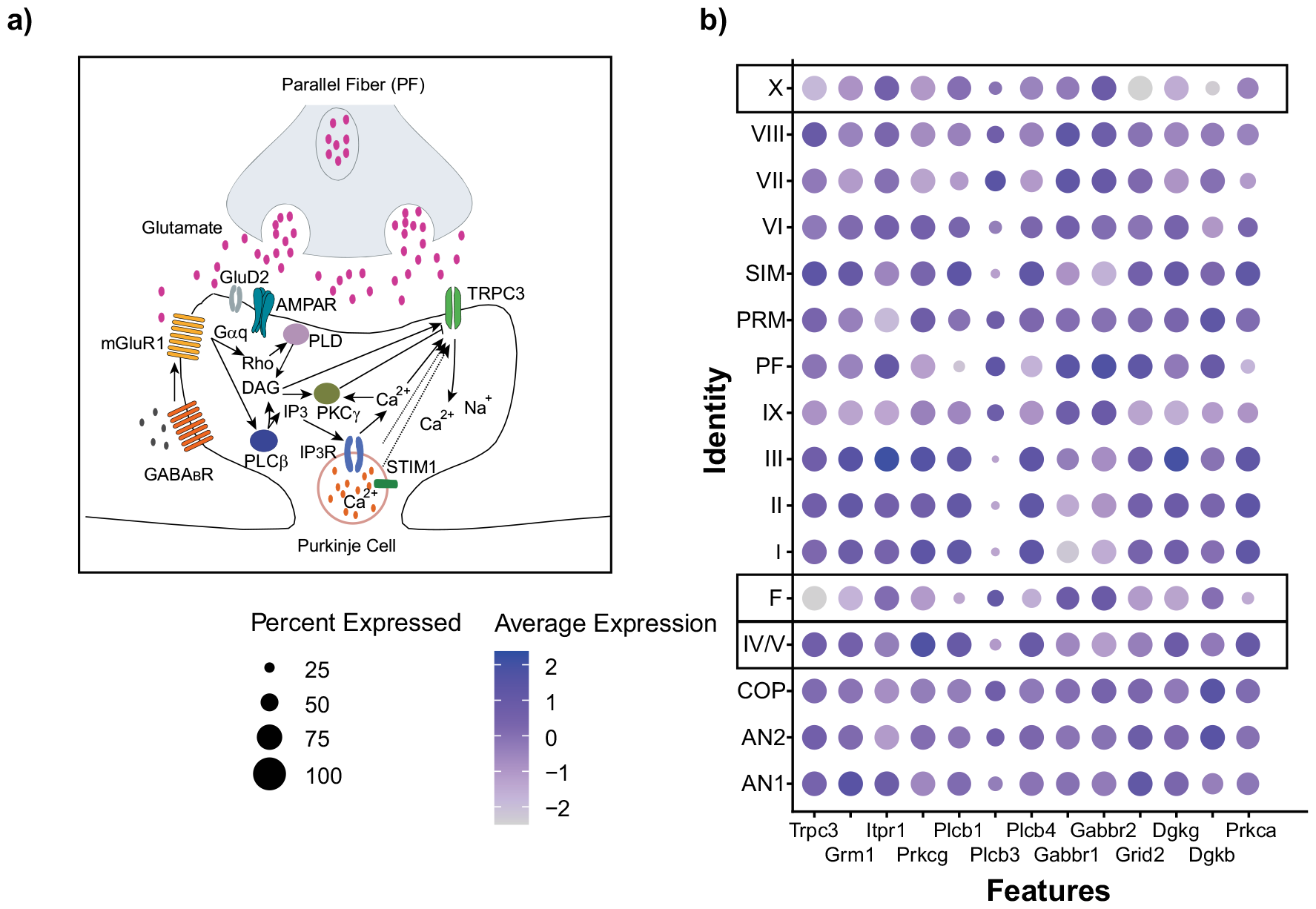
Heterogeneity in mGluR1-TRPC3 signaling in lobule IV/V, lobule X and the. a) Schematic of postsynaptic signaling pathway involved in the slow EPSC at PF to Purkinje cell synapses, combined and adapted from (Brasnjo and Otis, 2001; Cole and Becker, 2023; Hartmann et al., 2011; Hirai and Kano, 2018; Kano and Watanabe, 2017). b) Visualization of gene expression heterogeneity in single nucleus RNA-seq data of 16,634 Purkinje cells, from Kozareva et al. 2021. Dot plot shows scaled expression of selected genes of interest across cerebellar regions. The regions of interest are highlighted with black boxes. The following transcripts code for the protein of interest in Figure 5a above: Trpc3: TRPC3, Grm1: mGluR1, Itpr1: IP3 receptor, Prkcg: PKCγ, Plcb1, 4: isoforms of PLCβ, Gabbr1, 2: GABA_B_ receptor, Grid2: GluD2 receptor, Dgkb, g: Diacylglycerol kinase isoforms, Prkca: protein kinase Cα.

We then focused our analysis on the molecules that distinguish regions with slow timing (lobule X and flocculus) from regions with fast timing (lobule IV/V). We found that there were several differences in gene expression, including in the following: 1) Prkcg, which encodes PKCγ; 2) Plcb1 and 4, encoding PLCβ; 3) Dgkg, encoding diacylglycerol (DAG) kinase γ; 4) Gabbr2, encoding GABA_B_ receptors. In addition, Grid, encoding GluD2 channels, had lower expression in lobule X relative to the other two regions (see Discussion). Thus, we identified more than one molecular correlate of heterogeneity across lobules, within known components of mGluR-TRPC3 signaling pathways. In addition, these signaling components are likely to interact with each other (Cole and Becker, 2023). Moreover, our bioinformatic analysis by separating cells from different lobules obscures the within-lobule heterogeneity observed both in previous studies of the Purkinje cell transcriptome and in our data (Kozareva *et al*., 2021). Overall, we found several key differences in the signaling pathways that could mediate slow EPSC heterogeneity across lobules.

Analyzing their single-cell RNA-seq dataset of cerebellar Purkinje cells, Kozareva et al. (Kozareva *et al*., 2021) describe 9 molecularly-defined clusters that could be separated by zebrin identity: 2 clusters of zebrin-negative cells, and 7 of zebrin-positive cells. We examined the transcriptomic profiles of Purkinje cells from the three lobules that we are interested in, in terms of these 9 clusters (Figure S5). Surprisingly, lobular clusters do not fully align with the clusters defined by gene expression pattern. This observation may serve to explain the large amount of heterogeneity that we found across single cells. It also suggests that there is high-dimensional transcriptomic heterogeneity within and across regions of the cerebellum.

Next, to confirm our findings from a published RNA-seq dataset and to further investigate the receptor and channel mediating the slow EPSC, we performed quantitative reverse-transcription PCR analysis. We separated lobule IV/V and X of the vermis, as well as the flocculus, extracted RNA, and tested the relative abundance of mGluR1, TRPC3, and their splice variants (Figure 6a-f). Lobule IV/V showed higher expression levels of mGluR1b when compared to lobule X and the flocculus, which agrees with published differences in the isoforms of mGluR1 across zebrin-positive vs. negative regions (Mateos *et al*., 2001). Surprisingly, we also found a difference between the expression of the recently described splice variants of TRPC3, TRPC3c and 3b (Y. Kim *et al*., 2012). TRPC3c showed increased expression in lobule IV/V, the region with fast kinetics, compared to lobule X and the flocculus. In agreement with previous literature showing an anti-correlation between the expression levels of TRPC3 and zebrin across lobules (Wu *et al*., 2019), the expression of total TRPC3 was also higher in lobule IV/V. Taken together, our results identified a correlation between the expression levels of the splice variants TRPC3c and mGluR1b, and shorter vs. longer dynamics of the slow EPSC.

**Figure 6.**
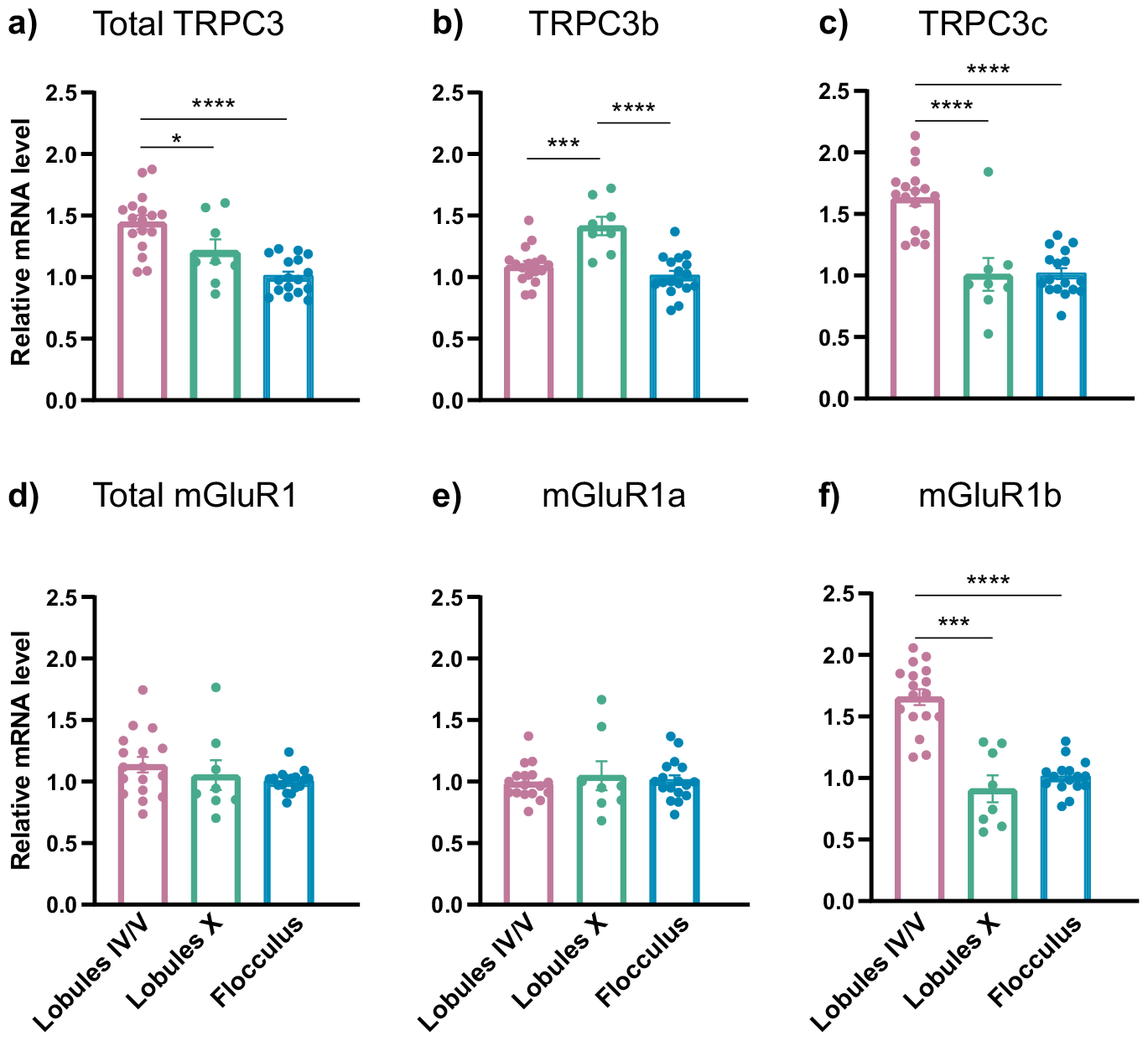
TRPC3 and mGluR1 isoform expression was heterogeneous across cerebellar lobules. Relative mRNA levels of a) total TRPC3, b) TRPC3b isoform, c) TRPC3c isoform, d) total mGluR1, e) mGluR1a, f) mGluR1b. (a,b,c) *p<0.05, ***p<0.001, ****p<0.0001, ordinary one-way ANOVA followed by Tukey’s multiple comparison test. (d,e,f) ***p<0.001, ****p<0.0001, Kruskal-Wallis test followed by Dunn’s multiple comparison test. All data are Mean ± SEM.

### Slow EPSC heterogeneity diversified timing of Purkinje cell firing

Next, we wondered how the slow EPSC heterogeneity would impact Purkinje cell spiking. It is known that the firing properties of Purkinje cells in response to depolarization are not uniform across the cerebellum (C. H. Kim *et al*., 2012; Zhou *et al*., 2014). To explore this, cells were first recorded in voltage-clamp configuration to measure the slow EPSC (Figure 7a-c). Next, recordings were switched to current clamp configuration, allowing Purkinje cells to fire action potentials in response to synaptic input (Figure 7d-f). This allowed us to look directly at any relationship between slow EPSC timing and parallel-fiber evoked modulation of spiking. We uncovered a difference in the timing of the firing response across lobules that matched the slow EPSC kinetics. Lobule IV/V, which had faster EPSC kinetics, had an equivalently short firing response. In contrast, slow EPSCs in lobule X and the flocculus, which had a diversity of longer delays, triggered prolonged firing responses in comparison to the responses elicited in lobule IV/V (Figure 7g-i).

**Figure 7.**
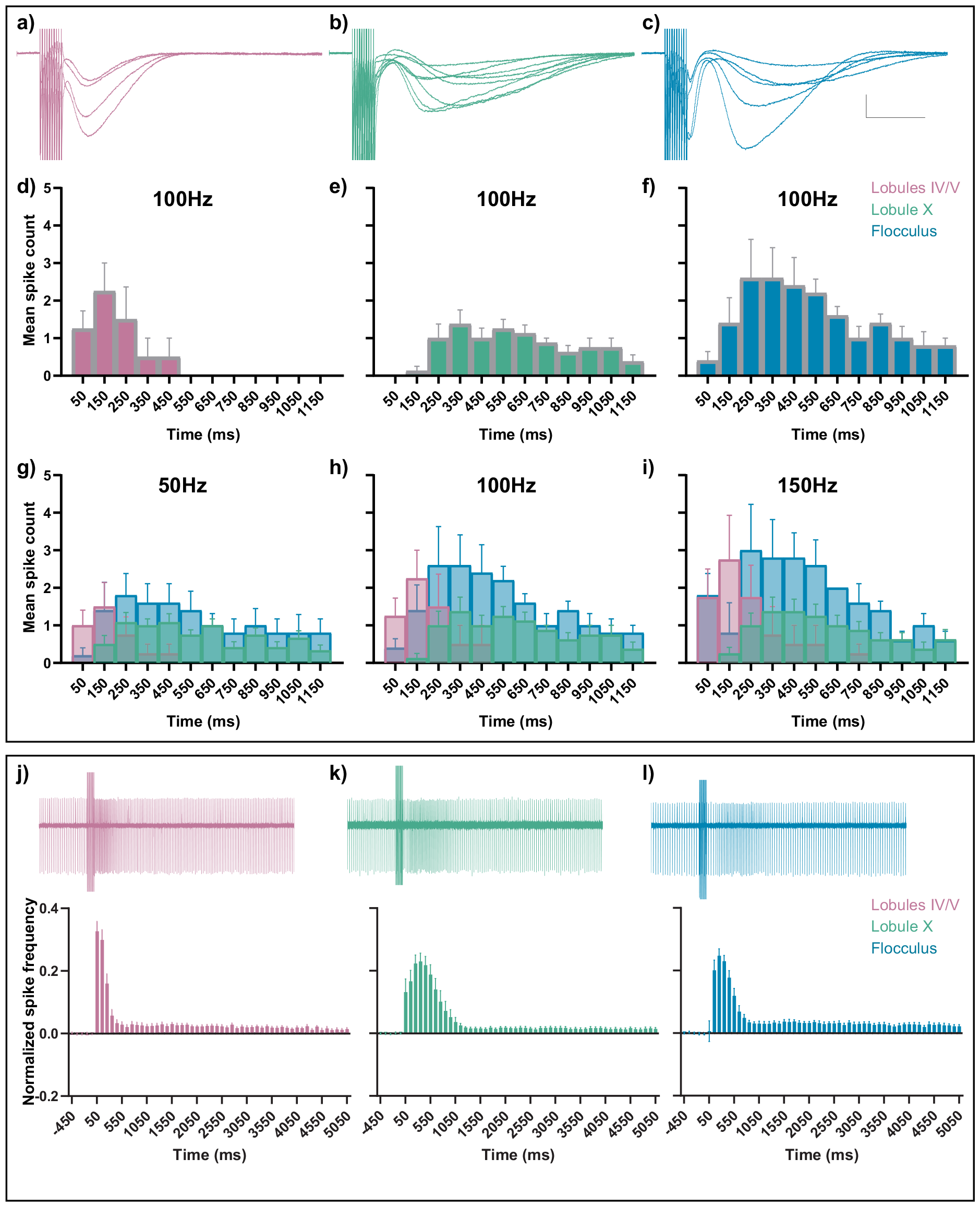
sEPSC heterogeneity diversifies Purkinje cell spike timing. (a, b, c) sEPSC traces in response to 10 PF stimulations at 100 Hz recorded from Purkinje cells voltage clamped at -70mV. Scale bar 100 pA, 250 ms (d, e, f) Mean spike counts binned into 100ms time bins from the same Purkinje cells, spiking in current clamp conditions, in response to the same stimulation. (g, h, i) Overlay of 100ms time binned mean spike counts from the same Purkinje cells under current clamp conditions in response to 10PF stimulations at (g) 50Hz, (h)100Hz, (i) 150Hz. (j) Top: Representative trace of a cell-attached recordings from lobule IV/V, Bottom: Normalized instantaneous spike frequency binned into 100ms time bins. The representative cell-attached trace is aligned in time to the x-axis of the graph. (k) Same as in figure 7j but for lobule X. (l) Same as in figures 7j,7k but for the flocculus. All data are Mean ± SEM.

We were concerned that the potentially dialyzing effects of whole cell recordings (Sakmann and Neher, 1984; Rae and Fernandez, 1988; Cahalan and Neher, 1992) affected our results. We also wondered what the effect of slow EPSC heterogeneity was on intrinsically firing cells. To address these concerns, we explored how slow EPSC heterogeneity affected Purkinje cell firing using cell-attached recordings. To isolate slow EPSC-driven effects, both AMPARs and GABA_A_ receptors were blocked. We observed that different regions of the cerebellum showed heterogeneity in the timing of their firing response (Figure 7j-l), which validated our previous whole-cell recordings (Figure 7a-i). The enhancement of firing was brief in lobule IV/V, in comparison to the delayed and prolonged enhancement of firing in lobule X and the flocculus.

Taken together, our findings demonstrate that the same synaptic input led to an increase in firing rate over different timescales across lobules (Figure 7j-l). Thus, the same synaptic input diversified PF-triggered responses heterogeneously across the cerebellum.

## Discussion

Our study reveals how Purkinje cells in different locations of the cerebellum transform the same synaptic input into heterogeneous outputs that vary in timing and dynamics. The difference in slow EPSC timing across lobules of the cerebellum correlated well with components of the mGluR-TRPC3 signaling pathway, implicating multiple molecular players in the regulation of slow synaptic signaling. Our results highlight a hitherto unappreciated form of cellular heterogeneity that complements and expands the known repertoire of cerebellar diversity (Ishii *et al*., 2010; Cerminara *et al*., 2015).

### The role of zebrin

We assessed the relationship of our observations of Purkinje cell heterogeneity in light of zebrin-positive or zebrin-negative identity. However, rather than having just two possible types of molecular signature, it has recently been described that Purkinje cells have diverse molecular properties (Guo *et al*., 2021; Kozareva *et al*., 2021). In addition, zebrin expression is also not necessarily binary, i.e., solely zebrin-positive or zebrin-negative, but has been described as having graded expression (Fujita *et al*., 2014). A broad diversity of cellular properties would serve to expand the computational capacity of the cerebellum. Our analysis of Purkinje cell transcriptomes also supports this idea of diversity in multiple different interacting molecules.

As a consequence of focusing on the slow EPSC, our study highlights the mGluR pathway (Reiner and Levitz, 2018). mGluR1 and its downstream signaling cascade is essential for normal cerebellar function, and mutations in mGluR1 impair motor coordination, cerebellum-dependent learning, and proper circuit development (Aiba *et al*., 1994; Ichise *et al*., 2000; Ohtani *et al*., 2014). Both mGluR signaling and TRPC3 signaling are critical for cerebellar synapses to function normally (Yamakawa and Hirano, 1999; Kishimoto *et al*., 2002; Knöpfel and Grandes, 2002; Hartmann *et al*., 2004, 2008; Lüscher and Huber, 2010; Hartmann and Konnerth, 2015; Kano and Watanabe, 2017). They are also important to understand as therapeutic targets because of their dysfunction in cerebellum-dependent diseases (Knöpfel and Grandes, 2002; Becker *et al*., 2009; Lüscher and Huber, 2010; Becker, 2014; Dulneva *et al*., 2015; Meera, Pulst and Otis, 2017; Tiapko and Groschner, 2018; Crupi, Impellizzeri and Cuzzocrea, 2019; Cole and Becker, 2023). Zebrin was potentially a relevant marker of heterogeneity for our study because of zebrin-based patterning of components of the mGluR-signaling pathway (Baron *et al*., 1999; Mateos *et al*., 2001; Wadiche and Jahr, 2005; Sarna *et al*., 2006; Furutama *et al*., 2010; Y. Kim *et al*., 2012; Wu *et al*., 2019; Cole and Becker, 2023). However, our data demonstrated that zebrin-identity alone did not determine slow EPSC timing.

### Key molecular determinants of synaptic diversity

Our analysis of published Purkinje cell transcriptomes to investigate regional differences highlights several molecules that are differentially expressed in the lobules with different slow EPSC dynamics. There is an increase in Prkcg expression, which encodes PKCγ, in the region with faster slow EPSCs relative to the other two regions. As summarized in Figure 5a, PKCγ negatively regulates TRPC3, which would be consistent with a shorter or more tightly timed slow EPSC (Hartmann, Henning and Konnerth, 2011; Cole and Becker, 2023). Plcb1 and 4, encoding PLCβ (Hashimoto *et al*., 2001; Miyata *et al*., 2001), are also differentially expressed in lobule IV/V vs. lobule X and the flocculus. Potentially, this could lead to faster activation of TRPC3 in lobule IV/V. In addition, Dgkg, encoding diacylglycerol (DAG) kinase gamma, is also higher in lobule IV/V. This is particularly interesting, as DAG-kinase has been shown to directly control the response duration of mGluR1 (Guo *et al*., 2021), which is consistent with our results. Grid2, encoding GluD2 channels, had a lower expression in lobule X relative to the other two regions. Since loss of GluD2 regulates mGluR-TRPC3 signaling (Kato *et al*., 2012; Ady *et al*., 2014), this may contribute to the characteristically small-amplitude and long-decay EPSCs in lobule X. Our analysis also showed that Gabbr2, encoding GABA_B_ receptors, was expressed at a higher level in the slow dynamics regions, i.e., lobule X and the flocculus, relative to the fast dynamics region, lobule IV/V. GABA_B_ receptors have been implicated in mGluR1-TRPC3 interactions and in the mGluR1-dependent slow EPSC, although the mechanisms by which it does so remain unknown (Tian and Zhu, 2018). Our comparison of lobular identity with previously described, transcriptionally-defined categories of Purkinje cells also suggests a diversity of categories within a lobule (Figure S5) (Kozareva *et al*., 2021). Moreover, Kozareva et al. described greater heterogeneity in zebrin-positive clusters of Purkinje cells (Kozareva *et al*., 2021), an observation that correlates with the greater heterogeneity in slow EPSC properties that we see in the largely zebrin positive lobule X and flocculus, in comparison to the relatively uniform slow EPSC properties in lobule IV/V. Our findings are also consistent with a recent study (Guo *et al*., 2021), which demonstrated diversification of timing in a cell-autonomous manner by mGluR1 signaling, in the context of cerebellar unipolar brush cells.

### Heterogeneity of TRPC3 isoforms

Our results also demonstrated a correlation between increased expression of the splice variant TRPC3c and shorter slow EPSCs in lobule IV/V. TRPC3c is distinguished from TRPC3b by the loss of a single exon, which codes for a large part of the calmodulin-inositol triphosphate (IP3) receptor binding domain (Y. Kim *et al*., 2012). Thus, it has altered intracellular regulation and sensitivity to Ca^2+^. In addition, there is an increased channel opening rate. However, the regulation and function of TRPC3 splice variants in Purkinje cells *in vivo* is not well understood, and it remains unclear how TRPC3c alone can directly cause shorter slow currents (Hartmann, Henning and Konnerth, 2011; Nelson and Glitsch, 2012; Y. Kim *et al*., 2012; Hartmann and Konnerth, 2015; Sierra-Valdez *et al*., 2018). Indeed, our results also demonstrate that there are additional molecular players that are likely to be responsible for different shape and timing of the slow EPSC. Overall, our findings point to the necessity to more fully understand expression and regulation of the mGluR-TRPC3 cascade (Bodzęta, Scheefhals and MacGillavry, 2021; Cole and Becker, 2023), and that previously-undescribed regional heterogeneity in splice variants needs to be considered.

### Implications for plasticity

Both mGluR1 and TRPC3 are essential for long-term depression (LTD) at PF to Purkinje cell synapses (Aiba *et al*., 1994; Chae *et al*., 2012; Kim, 2013), and LTD is critical for behavioral learning (Suvrathan, Payne and Raymond, 2016; Suvrathan and Raymond, 2018; De Zeeuw, Lisberger and Raymond, 2021). However, LTD is directly dependent on the postsynaptic Ca^2+^ signal (Wang, Denk and Häusser, 2000; Ito, 2001). The slow EPSC and the Ca^2+^ signal are thought to be two independent intracellular signaling cascades downstream of mGluR1 (Takechi, Eilers and Konnerth, 1998; Hartmann, Henning and Konnerth, 2011; Hartmann and Konnerth, 2015; Ouares and Canepari, 2020). Therefore, the timing of the slow EPSC is likely not related to the timing requirements for plasticity. However, it remains unclear how the diverse response timings of TRPC3 interact with heterogeneous plasticity mechanisms across the cerebellum, such as the demarcation into upbound and downbound zones (De Zeeuw, 2021).

### The link to Purkinje cell firing patterns

The slow EPSC-driven heterogeneity in firing we describe here adds to previous descriptions of differences in firing rate between Purkinje cells in zebrin-positive and zebrin-negative regions (Zhou *et al*., 2014), and the diverse intrinsic input-output relationships in different lobules (C. H. Kim *et al*., 2012). It was previously known that mGluR1b blockade can reduce Purkinje cell firing frequency (Yamakawa and Hirano, 1999), and TRPC3 blockade can affect Purkinje cell firing rate in a zebrin-dependent manner (Sekerková *et al*., 2013; Zhou *et al*., 2014), although the underlying mechanisms remain unclear.

The high-frequency spontaneous firing of Purkinje cells encodes the output of the cerebellar cortex and provides tonic inhibition to downstream cells in the deep cerebellar and vestibular nuclei. Consequently, the impact of homogeneous or diversified slow current timings, such as we observe in lobule IV/V vs. lobule X and the flocculus, is that it could affect the synchrony or lack thereof of Purkinje cells that are activated by the same PF beam. As a result, time-locked spiking of downstream deep cerebellar nucleus (DCN) neurons may be affected, as well as a direct impact on the firing rate of DCN neurons (Telgkamp and Raman, 2002; Person and Raman, 2012; Sedaghat-Nejad *et al*., 2022).

### Future directions

Our results highlight the importance of synaptic heterogeneity in Purkinje cell timing, which opens new areas of enquiry into how cerebellar circuits utilize this diversity. Our findings also expand the repertoire of synaptic and cellular mechanisms that different regions of the cerebellum may draw on to support diverse functions. Finally, although we did not causally link mGluR-TRPC3 signaling to this diversity, we reveal tight and suggestive correlations with key molecular players, which provides a firm foundation for future mechanistic studies into the role of synaptic heterogeneity in Purkinje cell timing. Our study is thus not a final verdict on Purkinje cell timing diversity, but rather a key starting point.

## Acknowledgements

We thank Dr. Jesper Sjöström, Dr. Keith Murai, Dr. Alanna Watt, and Dr. Arnold Hayer for their thoughtful comments and suggestions. We also thank Dmitri Yang and Chloe Guo for early experiments standardizing antibody staining protocols. R.E.T. was supported by an RIMUHC studentship and an IPN studentship, F.M. was supported by an RIMUHC studentship, A.S. received funding from CIHR Project Grant 178281, NSERC Discovery Grant, CFI-JELF, FRQS Chercheurs Boursiers/ Chercheuses Boursières, FRQS Établissement de Jeunes chercheurs, and startup funding from the Research Institute of the McGill University Health Centre and CFREF-HBHL. This paper was typeset with the bioRxiv word template by @Chrelli: www.github.com/chrelli/bioRxiv-word-template

## Author contributions

R.E.T and A.S. planned the experiments, R.E.T, F.M, K.S., and W.T.F. performed and analyzed the experiments, W.T.F. analyzed the bioinformatics, A.S., R.E.T. and F.M. wrote the paper.

## Materials and Methods

All animal experiments were done in accordance with the policies of the Canadian Council on Animal Care, using protocols approved by the Montreal General Hospital Facility Animal Care Committee.

### Ex vivo slice electrophysiology

C57Bl/6J mice (21-40 days) of both sexes from Jackson Laboratories (Strain number 000664) were used for all slice electrophysiology experiments. The cerebellum was dissected and 300 μm thick acute cerebellar slices were prepared in the sagittal (for vermis recordings) or coronal orientation (for flocculus recordings) using a Leica VT1200S vibratome in ice-cold artificial cerebrospinal fluid (aCSF) (concentration in mM): NaCl(119), KCl(2.5), NaH_2_PO_4_(1), NaHCO_3_(26.2), MgCl_2_(1.3), CaCl_2_(2.5), D-Glucose(10), equilibrated with Carbogen (95% O_2_, 5% CO_2_, Linde Canada)) or in ice cold sucrose cutting solution (concentration in mM): sucrose(200), KCl(2.5), NaH_2_PO_4_ (1), NaHCO_3_(26.2), MgCl_2_(1.3), CaCl_2_(2.5), D-Glucose(20). The slices were allowed to recover at 35 °C for 15-25 min in aCSF with constant carbogen bubbling, and then at room temperature for 1 hour.

Purkinje cells were visualized under an Olympus BX61WI upright microscope using differential interference contrast optics. Patch electrodes (3-6 MΩ) were pulled from borosilicate glass and filled with internal solution containing either one of the two recipes for both voltage and current clamp experiments (concentration in mM): Recipe1: Potassium gluconate(135), NaCl(7), MgATP(2), NaGTP(0.3), Hepes(10), EGTA(0.5), phosphocreatine di(tris) salt(10) (pH 7.2) Recipe 2: Potassium gluconate(128), KCl(4), Hepes(10), Sodium creatine phosphate(10), MgATP(4), NaGTP(0.3) (pH 7.3). For cell-attached recordings, the recording electrode was filled with NaCl (162.5mM) solution (Perkins, 2006). Stimulation of the parallel fibers was performed using bipolar stimulating electrodes made from theta glass that were positioned in the outer third of the molecular layer. aCSF contained 50 μM picrotoxin and 5μM NBQX for all recordings (except for in Figure S1, where only picrotoxin was added). Additional details can be found in Supplementary Methods.

### Fixed Tissue Preparation and Immunofluorescence

50 μm slice staining: Mice brains were fixed with 4% paraformaldehyde (PFA) using intracardiac perfusion followed by overnight fixation in PFA at 4° C. The cerebellum was sliced in the parasagittal or coronal orientation to obtain 50 μm slices using a Vibratome 1000 Plus. Slices were incubated for two days with primary antibodies. On the 3^rd^ day the primary antibodies were washed off and the section incubated in secondary antibodies for 90 to 120 minutes. Blocking with Fab fragments were performed when using mouse primary antibody. 300 μm slice staining: For immunostaining of 300 μm thick slices, post-electrophysiology and dye-fill (Figure 2e, f, g), the slices were fixed in PFA for 1 to 4 hrs. The immunostaining protocol was the same as for 50 μm slice staining except for a 3-day incubation time with primary antibody.

### Quantitative reverse transcription polymerase chain reaction (RT-qPCR)

The cerebellum was extracted, lobules of interest (lobules IV/V, lobule X and flocculus) dissected using a dissection microscope and snap frozen. Each sample was prepared by clubbing the lobule of interest from 3 animals. This was followed by RNA extraction and reverse transcription into cDNA. qPCR was performed on a StepOnePlus Real-Time PCR System (Applied Biosystems by ThermoFisher Scientific) using Fast SYBR Green Master Mix. Relative expression levels of each target transcript were determined by the ΔΔC? method (Livak and Schmittgen, 2001) using RPL13 mRNA levels as a reference.

### Single nucleus RNA-seq Analysis

The processed single cell RNA-seq data from Kozareva et al, 2021 was filtered to contain only the 16634 Purkinje cells using Seurat 5.0.1 in R 4.3.2. The slots containing metadata, raw counts, normalized, and scaled data were copied from the author’s Seurat 2.3.4 object found here: https://singlecell.broadinstitute.org/single_cell/study/SCP795/a-tran-scriptomic-atlas-of-the-mouse-cerebellum to create a new Seurat v5 compatible object. Variable features, principal component, and UMAP dimension reductions were then recalculated using the respective Seurat functions on the Purkinje-only data. Plots were produced using the DimPlot and DotPlot functions of Seurat and ggplot2. Differential gene expression analysis was performed using the Findmarkers function of Seurat. Lobules of interest were compared pairwise using the default Wilcoxon Rank Sum Test while considering all detected genes.

Additional details can be found in the Supplementary Methods.

