## Supplementary Information for "Heterogeneity in slow synaptic transmission diversifies Purkinje cell timing"

bioRxiv

**Supplementary Information**

### Figure S1

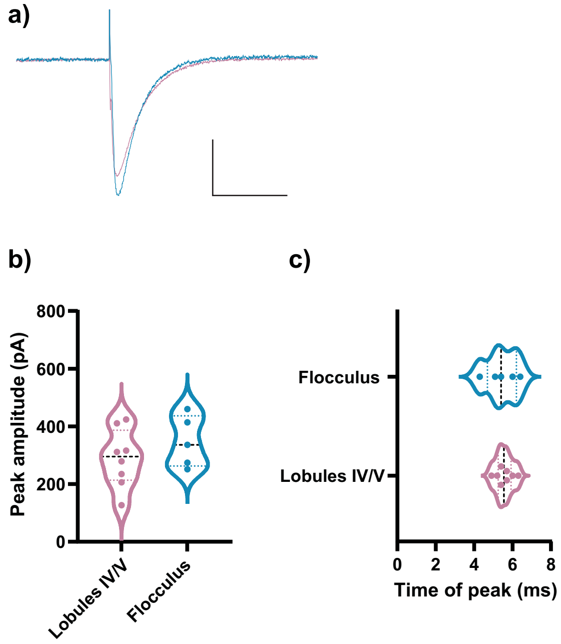


**Figure S1: AMPAR-mediated synaptic responses in lobule IV/V and flocculus were indistinguishable**

a) Representative traces of AMPAR-mediated EPSC from lobule IV/V and the flocculus. Scale bar 100 pA, 50 ms. Peak amplitude (b) and time of peak (c) were the same in lobule IV/V and the flocculus, in contrast to the time of peak of sEPSCs. Violin plots show median and quartiles.

**Figure S2**

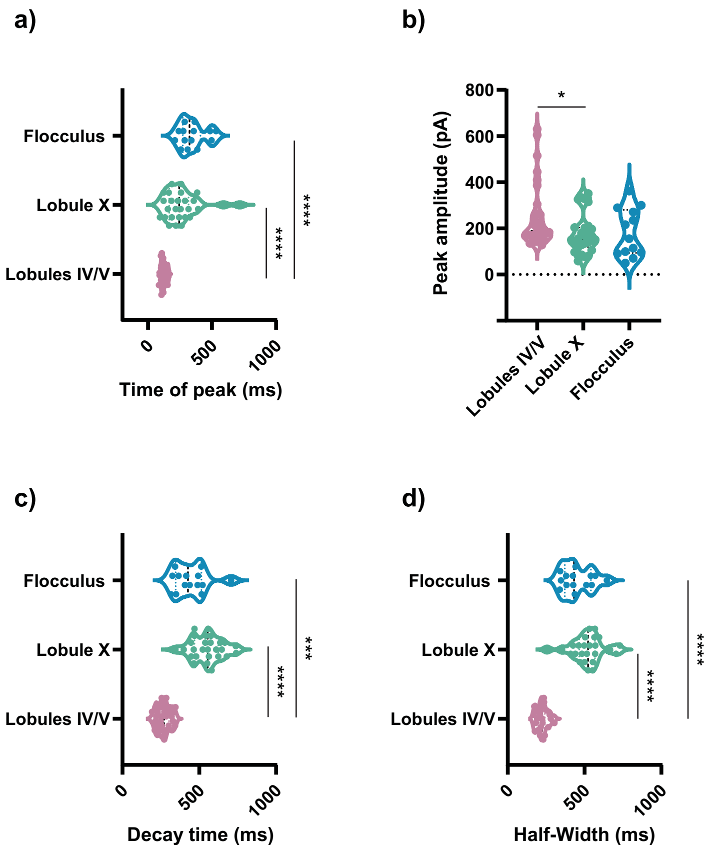


**Figure S2: Lobule-specific synaptic dynamics were not sensitive to Purkinje cell dialysis**
a) Time of peak of a subset of sEPSCs from Figure 1, across lobule IV/V, lobule X and the flocculus, recorded within 10-20 min after break-in to whole-cell configuration, demonstrated shorter time of peak in lobule IV/V.
b) Peak amplitude of subset of sEPSCs was similar across regions
c) Decay time of sEPSCs were shorter in lobule IV/V, in comparison to lobule X and the flocculus, even for this subset of cells.
d) Half-widths of sEPSCs were shorter in lobule IV/V, in comparison to lobule X and the flocculus, even for this subset of cells.
Statistical comparisons: (a,b,c,d)*p<0.05, , ***p<0.001, ****p<0.0001, Kruskal-Wallis test followed by Dunn's. Violin plots show median and quartiles.

### Figure S3

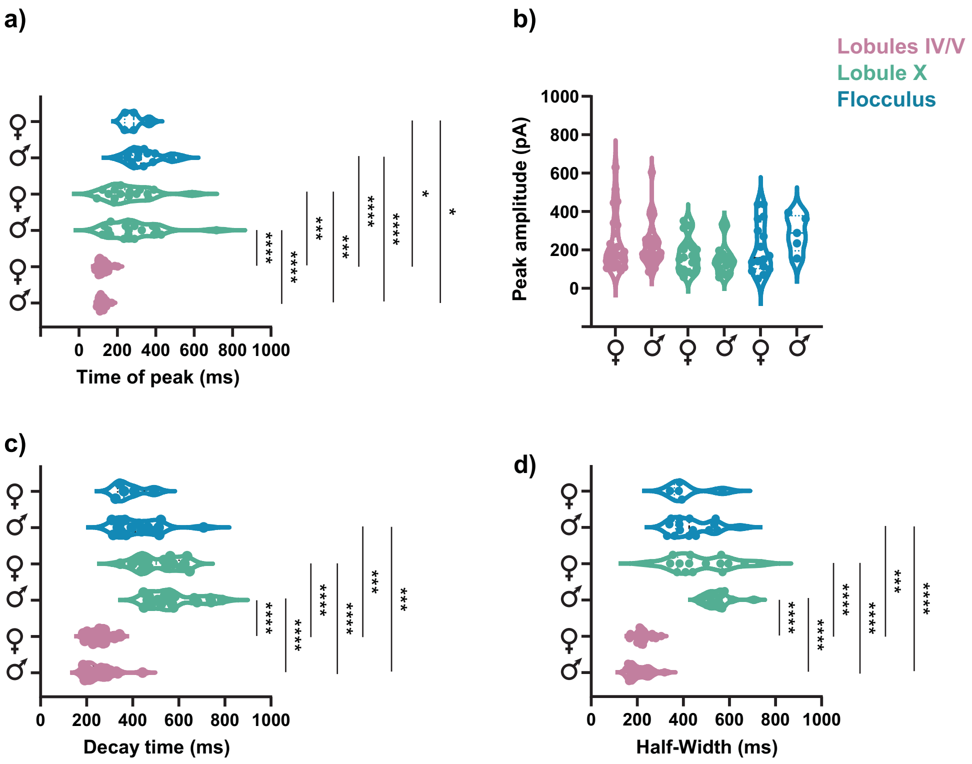


**Figure S3. Animal sex did not determine lobule-specific synaptic dynamics**
a) Subset analysis of data in Figure 1 demonstrated that the sEPSC time of peak did not differ across sexes, nor did peak amplitude (b), decay time (c), or half-width (d).
Statistical comparisons: (a,b,c,d) *p<0.05, ***p<0.001, ****p<0.0001, Kruskal-Wallis test followed by Dunn's multiple comparison test.

**Figure S4**

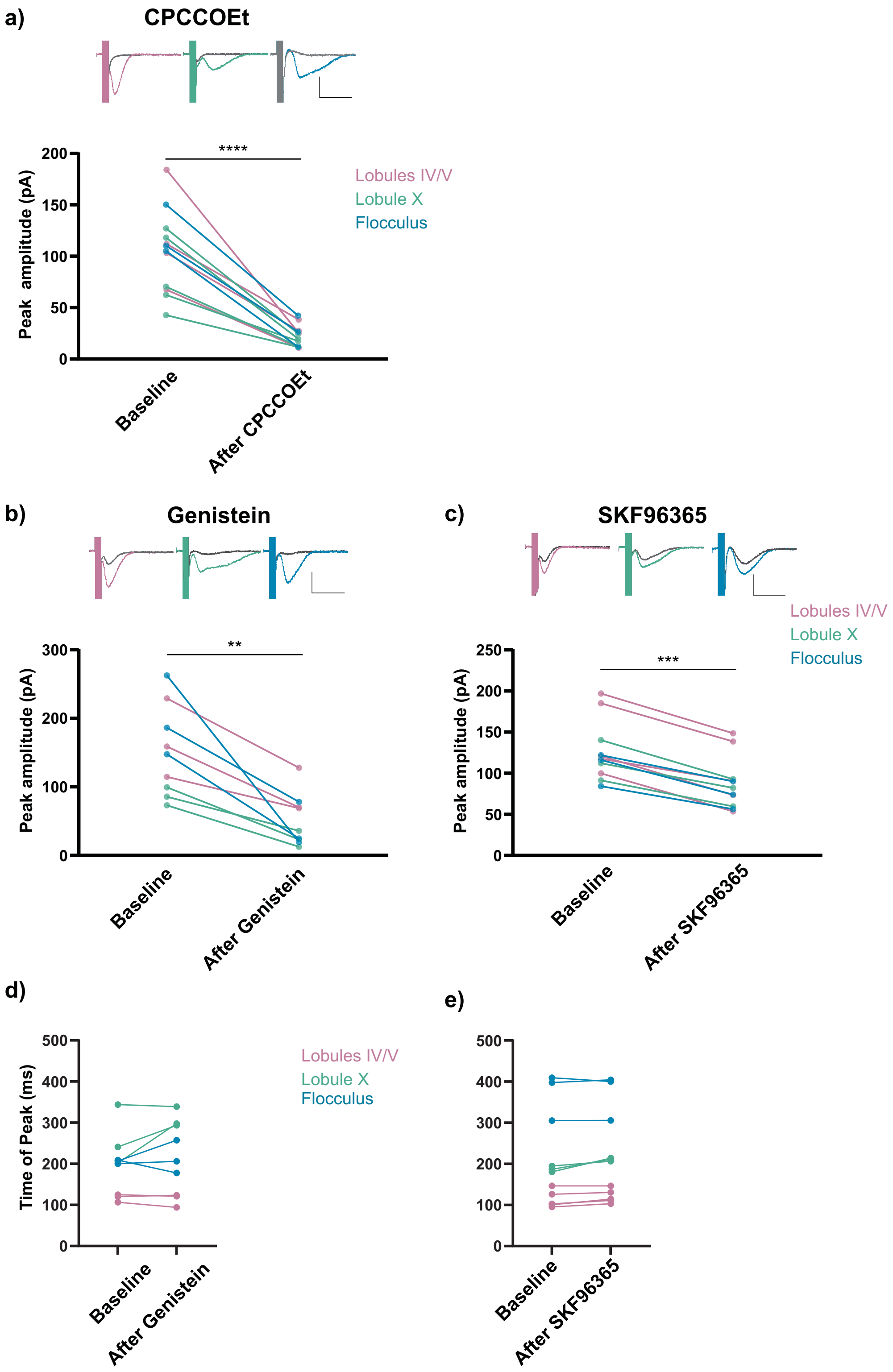


**Figure S4. sEPSCs exhibited equivalent mGluR- and TRPC3-dependence across lobules**
a) Top: Representative sEPSC traces from lobule IV/V, lobule X and flocculus. Colored traces are sEPSCs at baseline and grey traces represent sEPSCs after blockade by wash-in of noncompetitive mGluR 1 antagonist CPCCOEt. Bottom: sEPSCs were blocked by wash-in of CPCCOEt. *p<0.05, paired t-test.
(b,c) Top: Representative sEPSC traces from lobule IV/V, lobule X and flocculus. Colored traces are sEPSCs at baseline and grey traces are sEPSCs after wash-in. Bottom: (b)Wash-in of Genistein (100 μM) or (c) SKF96365 (10 μM) reduced sEPSC amplitude across all cerebellar lobules tested. **p<0.01, ***p<0.001, Wilcoxon matched-pairs signed rank test.
(d, e) Partial blockade of TRPC3 did not explain heterogeneity in time of peak across lobules. Statistics were not run to compare different lobules as the number of cells in each lobule was too low.
Scale bars are 100 pA, 0.5 s. Violin plots show median and quartiles.

### Figure S5

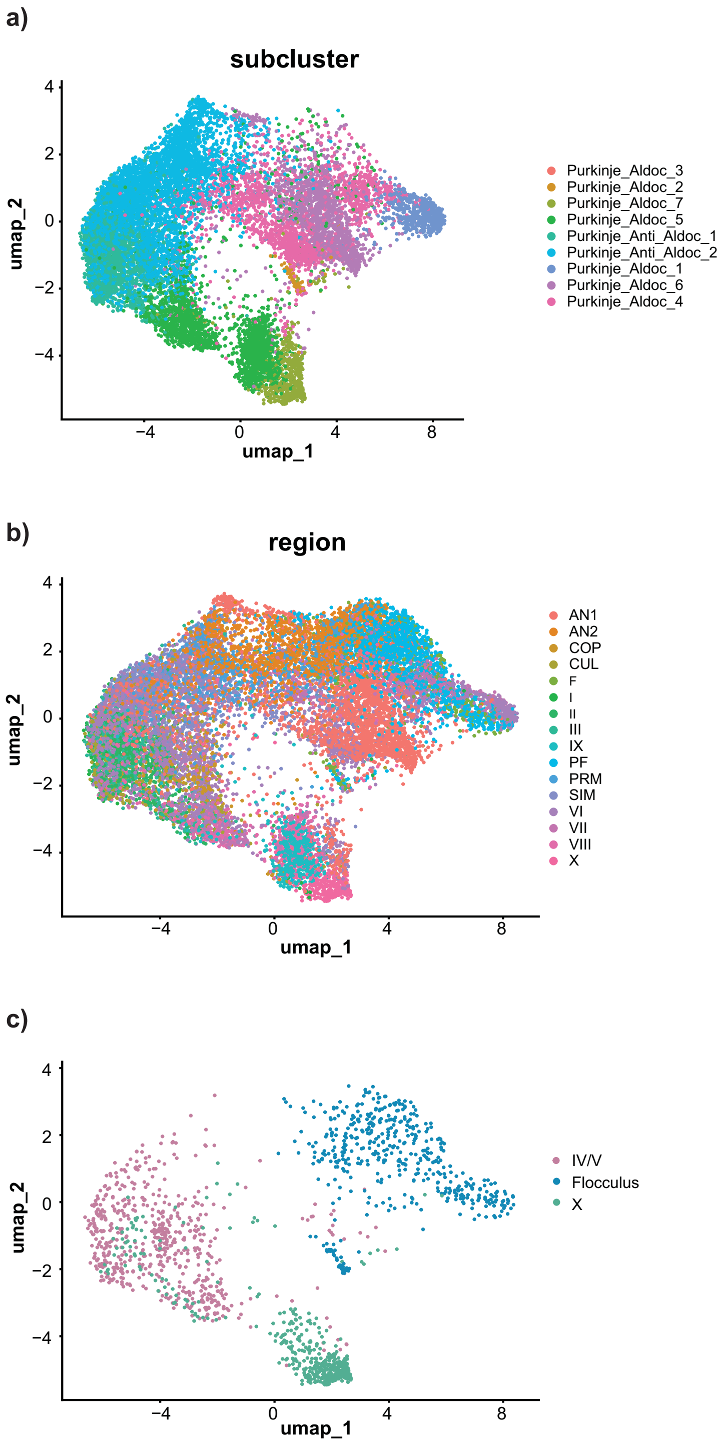


**Figure S5. Gene expression is heterogeneous across lobule IV/V, lobule X, and the flocculus**
Visualization of gene expression heterogeneity in scRNA-seq data of 16634 Purkinje cells from Kozareva et al. 2021. a) UMAP dimension reduction showing original authors' identified PC clusters. Aldoc and zebrin are equivalent b) UMAP plot showing the region of the cerebellum that the cells were sampled from, demonstrating the lack of full alignment between molecularly-defined cell-type and lobular identity. c) Lobule IV/V, lobule X and the flocculus were separated out from b), demonstrating that all three regions fall in different clusters, with some overlap.

### Supplementary Methods

Ex vivo slice electrophysiology: All whole cell patch clamp recordings were performed at 29-30°C. For cell attached recordings the temperature was maintained at 33±1°C. Signals were acquired using a MultiClamp 700B amplifier at 10-100 kHz for the slow EPSC and 50 kHz for spiking experiments. The slow EPSC traces were averaged and filtered using a Bessel (8-pole) filter with low pass filter of 1kHz. Only recordings with access resistance <25 MΩ were included. Both input and access resistance were monitored for stability, and discarded if they changed more than 20%. Slow EPSCs were elicited using 10 PF stimulation at a 100 Hz, with 100 µA stimulation current intensity for all experiments except figures 3 and 4, where the frequency and number of PF stimulation were varied, as described in the figure legend. For voltage clamp recordings the cells were held at -70mV. For the whole cell spiking experiments (Figure 7 a -i)) the slow EPSC traces were first recorded in voltage clamp configuration and consequently switched to current clamp with sufficient current injection to keep the cells below the threshold for spiking (between –49 to -57 mV at baseline, except in one flocculus cell where the cell was spiking at baseline even at this voltage). For figures 3 and 4 some cells that contributed to the data came from the same cells recorded in figure 1, figure 7 a-i, or from figures S4 a-e.

Cell-attached recordings were analyzed by normalizing the instantaneous firing frequency according to the following equation: (Inst. Freq. – Mean baseline Inst. Freq.)/ (Inst. Freq. + Mean baseline Inst. Freq.).

Reagents were obtained from the following vendors: NBQX (Sigma Aldrich N-183 or Abcam ab120046), CPCCOEt(Sigma Aldrich SML1124 or Abcam ab120060), Alexa Fluor 488 Hydrazide (Thermo Scientific A10436). NBQX (N183 ), Picrotoxin (P1675), Genistein (G6776), and SKF-96365 (S7809) were all obtained from Sigma Aldrich. TBOA (1223/10) was obtained from Tocris Bioscience.

Fixed Tissue Preparation and Immunofluorescence imaging:

Fab fragment used was AffiniPure Fab Fragment Donkey Anti-Mouse by Jackson Immunoresearch Labs. (715-007-003) at 1:50 or 1:200 dilution. Prior to mounting on a slide, slices were incubated in Hoechst 33342 (1:10000) when using mounting medium without DAPI.

For figure 2a, b, c, e, f, g blocking solution used contained 1X PBS, pH 7.4, 0.4% Triton X, 5% bovine serum albumin (BSA), and 0.05% sodium azide.

For Fig 2d, prior to staining, slices were subjected to an antigen retrieval procedure. Slices were boiled at 90°C in 10 mM tri-sodium citrate buffer (pH 8.5) for 10 min followed by a 20-minute cool-down period. The blocking solution used contained 1X PBS, pH 7.4, 0.4% Triton X, and 8% heat-inactivated normal goat serum.

Epifluorescence images were captured using a fully automated epifluorescence microscope system (TI2-E, Nikon), equipped with a multispectral LED light source (SpectraX, Lumencor) and 10x Plan Apo 0.45 NA objective, a motorized emission filter wheel (HS 1025, FLI), a motorized stage, an sCMOS camera (ORCA Fusion BT, Hamamatsu) and controlled by Nikon Elements AR. Images were acquired as multi-point images and stitched together using Nikon Elements AR's "large image" feature. 300 µm thick slices immunostained after patch-clamp recordings were imaged on Olympus FV-1000 laser scanning microscope, with a 20X lens and acquired using Fluoview software (Olympus). Post-patch-clamp recording zebrin labelling was quantified by taking a ratio of fluorescent intensity within the Purkinje cell to the background fluorescence in the neighboring granule cell layer using Fiji (ImageJ). Only cells with a ratio >2 were considered zebrin-positive. Those with a ratio < 1.7 were considered zebrin-negative, and those in between were considered uncertain and were therefore not used in the analysis shown in Figure 2.

Quantitative reverse transcription polymerase chain reaction (RT-qPCR):

26 to 33 days old mice of both sexes were used. RNA extraction was done using the RNeasy Mini Kit (Qiagen- 74104) following the manufacturer's instructions. RNA was then transcribed into cDNA using the QuantiTect Reverse Transcription Kit (Qiagen-205313). Fast SYBR Green Master Mix (Applied Biosystems-4385612) was used for the qPCR. qPCR reactions were run as technical triplicates in 10 µL reactions (96-well format).

The thermal cycling was performed with the following settings: Holding stage: 20 seconds at 95°C, cycling stage: 40 cycles of 3 seconds at 95 °C and 30 seconds at 62 °C. Melt curves were generated by heating from 62 °C to 95 °C at a temperature increment of 0.3°C.

Statistical analysis:

Slice physiology data was analyzed in pCLAMP 11 (Clampfit) and the statistical tests were performed using GraphPad Prism except for Levene's test which was performed on JASP statistics program.

For figure 1 c, d, f, g; figure 3 a-i; figure 4 a-c; figure 6a-f; figure S2 a-d and figure S3 a-d, the data was tested for normality using Shapiro-Wilk test and for equality of variances using Levene's test. If the data satisfied both the normality and equality of variances conditions, then it was analysed using ordinary one-way ANOVA followed by Tukey's multiple comparisons test. If either one of the assumptions was violated, Kruskal-Wallis test followed by Dunn's multiple comparisons test was used. For figure S4 a-c paired t-test was used if the data passed normality test using Shapiro-Wilk test. If not, Wilcoxon matched-pairs signed rank test was used. For figure S1 b, c unpaired t-test was used following normality check using the Shapiro-Wilk test.

| Primary antibodies | Dilution (For 50 µm slices) | Dilution (For 300 µm slices) | Vendor & Catalogue number | Figures |
|---|---|---|---|---|
| 1) Anti-Aldolase C antibody [4A9] - N-terminal | 1:300 | 1:100 | Abcam (ab190368) | 2 a-g |
| 2) Calbindin D28K rabbit recombinant polyclonal antibody | 1:500 | N/A | Thermo Scientific (PI711443) | 2 a, b, c |
| 3) Calbindin D28K chicken polyclonal antibody | 1:2000 | N/A | Thermo Scientific **(PA5143561)** | 2d |
| Secondary antibodies | Dilution (For 50 µm slices) | Dilution (For 300 µm slices) | Vendor & Catalogue number | Figures |
| 1) Goat anti-mouse Alexa Fluor 568 | 1:1000 | 1:1000 | Invitrogen (A11004) | 2 a, b, c, e, f, g |
| 2) Goat anti-rabbit Alexa Fluor 488 | 1:1000 | N/A | Invitrogen (A11008) | 2 a, b, c |
| 3) Goat anti-mouse Alexa Fluor 488 | 1:500 | N/A | Invitrogen (A11029) | 2d |
| 4) Goat anti-chicken Alexa Fluor 647 | 1:1000 | N/A | Invitrogen(A21449) | 2d |

| Primer name | Fwd (5'-3') | Rev (5'-3') |
|---|---|---|
| RPL13 | GGCCAGAGTTATCACAGAAGAA | TTGCTCGGATGCCAAAGA |
| Total TRPC3 | GATCGAGGATGACAGTGATGTAG | GACTGAAGGGTGGAGGTAATG |
| TRPC3b | TCCAAATGCAGGAGGAGAAG | CGAGTTAGACTGTGTGAAGAGG |
| TRPC3c | GTAACTCAAAGTCCAGGCAGATA | TCAGTTCACCTTCATTCACCTC |
| Total mGlu1 | TCTGTGATTCCCTGTGTTTAGG | CGTCTCGCTTCTCTCTTTACTC |
| mGlu1a | AGGGAACGCCAATTCTAACG | CCACACATGCTGTCCCTT |
| mGlu1b | CAAGAAGAGGCAGCCAGAA | GCCGTTAGAAACATTCACTGC |
